# LooMS: a novel peptide identification tools for data independent acquisition

**DOI:** 10.1101/2024.06.20.599973

**Authors:** Jiancheng Zhong, Jia Rong Wu, Xiangyuan Zeng, Michael Moran, Bin Ma

## Abstract

Advancements in mass spectrometry (MS)-based proteomics have produced large-scale datasets, necessitating the development of effective tools for peptide identification. Here, we present LooMS, a novel tool specifically designed for identifying peptides in data-independent acquisition (DIA) datasets. LooMS employs an innovative approach, using an unbiased generation strategy for positive and negative samples, which reduces the risk of overfitting in peptide identification with deep learning models. Additionally, LooMS addresses various critical aspects of DIA mass spectra data analysis, constructing a comprehensive set of 43 features for training deep learning models, which cover different stages of DIA data analysis. Notably, we propose a false discovery rate (FDR) control strategy that integrates results from both LooMS and DiaNN, another leading peptide identification tool. Our results demonstrate significant improvements in peptide identification performance, with enhancements of 40.61% and 26.60% at the unique peptide level for human and mouse datasets, respectively.

**Highlights:**

- LooMS is a novel tool for identifying peptides in DIA datasets that adopts an innovative unbiased positive and negative sample generation strategy, which aim to avoid the overfilling in peptide identification with deep learning model.
- LooMS comprehensively considers various aspects of data analysis for DIA mass spectra and builds 43 useful features for training deep learning models, which involve different stages of DIA data analysis.
- A FDR control strategy for integration of results from both LooMS and DiaNN is proposed, which can significantly improve the identification of peptides due to the differences in the features involved in peptide detection during their respective design.

## 1. Introduction

Biomarker identification, personalized medicine, and drug discovery have increased the demand for high-throughput proteomics[1]. To meet this demand, DIA mass spectrometry has been developed. Unlike traditional data-dependent acquisition (DDA) methods, DIA provides deeper proteome coverage in a single injection and at a faster rate. Moreover, DIA eliminates the need for manual intervention to select high-abundance peaks for further MS/MS analysis. This results in a more comprehensive and reproducible sample analysis, avoiding selection bias towards high-intensity precursors[2].

DIA achieves this by sequentially acquiring and analyzing all ions present in a sample, fragmenting all precursors at regular retention time intervals within user-configured mass spectrometry windows[3]. The window sizes can be adjusted to cover a few Daltons up to the entire mass range, allowing for a complete proteome profile acquisition.

The use of DIA approaches in mass spectrometry allows for the rapid generation of highly multiplexed tandem mass spectra. However, this presents significant computational challenges for peptide identification due to the extensive mixing of co-fragmenting precursors and co-eluting peaks within wide mass range windows. These challenges require novel computational strategies to address the issue of identifying individual peptides from such complex spectra.

Addressing the complexities of DIA mass spectrometry has been a major focus of research, leading to the development of several innovative strategies. These can be categorized into three main groups: spectrum-centric, peptidecentric, and hybrid methods.

Spectrum-centric methods such as DIA-Umpire[4] and Group-DIA[5] operate based on the principle of DDA database searching.. They generate pseudo MS/MS spectra to identify all possible precursor-fragment pairs based on their elution profiles.. However, these methods are unable to resolve undetectable precursors, a common occurrence in mass spectrometry.

Peptide-centric strategies, such as MSPLIT-DIA[6], PECAN[7], FT-ARM[8], and Diamond[9], adopt the theoretical spectra of peptides derived from reference protein sequences to interrogate DIA mass spectra. These methods offer higher sensitivity compared to spectrum-centric approaches and enhance peptide identification accuracy by incorporating retention time and fragment ion data from spectral libraries[10][11]. If spectral library is not available, traditional methods to generate spectral libraries often involve multiple DDA analyses, which can introduce bias and are time-consuming. These libraries are also limited to specific laboratories or instruments. To address this, in silico approaches, like Prosit[12], AutoRt[13], deepDIA[14], pDeep[15][16][17], and empirically corrected libraries[18], have been proposed to predict RT and fragment ion intensities, creating spectral libraries that improve the reproducibility and unbiased nature of DIA analysis.. Despite their high performance, peptide-centric approaches can struggle with peptides that have overlapping elution profiles or similar mass/charge ratios, potentially leading to incorrect identifications.

State-of-the-art methods for DIA analysis integrate the strengths of both peptide-centric and spectrum-centric approaches. Tools such as DiaNN[19], dia-PASEF[20],Alpha-Tri[21], MaxDIA[22], and MSFragger-DIA[23] implement feature detection techniques for precursor ions and their fragments while also generating in silico spectral libraries for database searches. These methods construct a feature space and apply machine learning algorithms to rescore peptide-spectrum matches (PSMs). However, training these models with various machine learning algorithms requires known samples containing target and decoy information, which can sometimes lead to overfitting.

Here, we introduce LooMS, a comprehensive tool for DIA data analysis that integrates the strengths of both peptide-centric and spectrum-centric approaches while addressing challenges at each analytical step. In the preprocessing stage, LooMS employs a trait tracing technique to identify highquality traits of precursor and fragment ions, incorporating co-eluting information. Additionally, it labels low-abundance precursors by examining their isotopes. During the database searching stage, LooMS generates pseudo-precursors to detect peptides with multiple charge states that lack an actual precursor peak signal. It also considers the entire retention time range of the precursor to trace fragment ions with similar elution profiles, including possible isotope trails of fragment ions. LooMS extracts a total of 43 features to create the feature space. To mitigate overfitting, LooMS employs a novel unbiased hidden-target strategy to generate positive and negative samples for modeling, using an iterative and boosting approach. A novel FDR control method is introduced to combine results from LooMS and DiaNN. The performance of LooMS was evaluated on two datasets, showing an improvement over DiaNN by 18.58% on the HELA dataset and 2.17% on the mouse dataset. Furthermore, combining the results from both methods enhanced performance by 40.61% on the HELA dataset and 26.60% on the mouse dataset compared to using DiaNN alone.

## 2. Experimental procedures

### 2.1. Overview of LooMS

The LooMS approach, illustrated in Figure 1, comprises three main components. Firstly, it conducts a thorough analysis of DIA mass spectrometry data at every stage. This includes processing raw signals, aligning peptide sequences with MS1 precursor and MS2 fragment ions, and integrating predictions from spectral libraries. LooMS constructs a comprehensive set of features for training deep learning models, covering the entire DIA data analysis workflow. Secondly, to mitigate overfitting, LooMS employs a novel sample partitioning model. This model generates positive samples by combining genuine target sequences with decoy sequences and forms negative samples by pairing two decoy sequences. By concealing the genuine target sequence, the model prevents direct access during training, thus reducing bias. This partitioning strategy enhances the generalizability of deep learning models in peptide spectrum matching. Lastly, LooMS uses an iterative approach to train deep models and applies bagging to combine multiple deep learning models, yielding the final PSM score. This comprehensive strategy effectively addresses DIA data analysis challenges, reduces bias, and improves the accuracy of PSMs identification.

**Figure 1:**
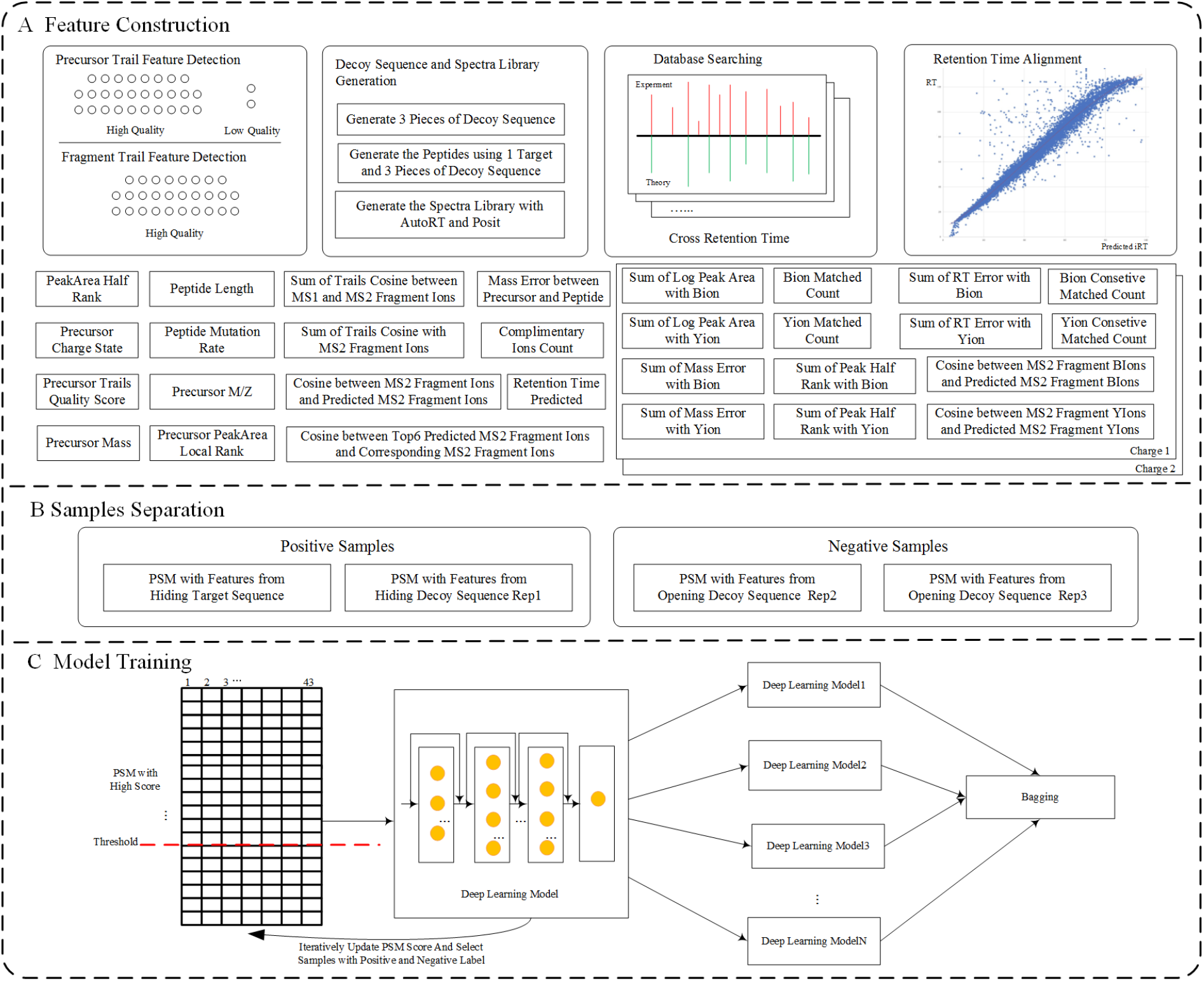
The pipeline of LooMS.

### 2.2. Data processing and feature space construction

#### 1. Extraction of Trails from Precursor and Fragment Ions

Elution profiles describe the behavior of a sample as it passes through a chromatography column, typically represented as a graph with retention time on the x-axis and peak intensity on the y-axis. LooMS uses various chromatogram extraction strategies to obtain trails from both precursor and fragment ions. For low-abundance analytes with poor ion signals, LooMS considers both high and low-quality signal trails. To address undetectable precursor signals due to intra-scan dynamic range limitations, LooMS generates pseudo-precursors for detecting peptides with different charge states when no precursor signal is present. For fragment ions, LooMS uses a stick strategy to select peaks, which avoids exceeding the dynamic range of MS/MS analysis. Additionally, to minimize interference from unfragmented precursors in the fragment ion chromatogram window, LooMS eliminates elution peaks within the precursor retention time window that have similar masses to the precursor. This preprocessing step enhances the accuracy of fragmentation spectra. The peak area of trails for each precursor and fragment ion is calculated by integrating ion intensity over the m/z range, serving as features in database searching. The shape of trails is also extracted to compare elution profiles and identify discrepancies. LooMS defines localHalfRank as a measure based on ion intensity within a specific m/z range (+/- 50 Da) and retention time window (+/- 5 seconds), ranking ions by peak area to identify more abundant ions in one group compared to another.

#### 2. Generation of Decoy Sequences and Spectral Library in Silicon

LooMS uses protein reference sequences to generate protein decoy sequences for FDR analysis and unbiased sample partitioning during model training. During protein digestion, peptides with missed cleavage sites can be produced. To account for this, LooMS includes a default parameter for missed cleavages, enabling the identification of such peptides. LooMS leverages cutting-edge spectral library and retention time prediction tools like Prosit and AutoRT to generate spectral libraries in silico, facilitating peptide detection. The default model and parameters of these tools help LooMS create a spectral library containing peptides of interest. These tools also contribute to database searching and feature engineering. For instance, a cosine similarity score is calculated to assess the similarity between candidate fragment peaks measured in database searches and their corresponding peaks predicted by Prosit. Despite data fluctuations from mass spectrometers, there is a strong correlation between predicted and measured retention times for high-scoring peptides. LooMS employs a retention time alignment algorithm to map predicted retention times to experimental retention times, controlling false matches in database searching.

#### 3. Database Searching for PSMs

After extracting precursors and fragment ions, LooMS performs spectral matching of peptides from both target and decoy sequences, generating initial PSM scores by identifying peptides that match the precursor mass and have more than three corresponding theoretical ions in the fragment ion spectrum. In DIA datasets, mixed precursors within an isolation window and multiplexed fragment ions pose challenges for peptide detection. To avoid erroneous matches with noise and interferent peaks, LooMS filters isotope peaks before database searching. It detects isotope peaks with different charge states and filters peaks meeting isotope abundance conditions, preventing them from participating in subsequent searches. LooMS performs database searches by detecting fragment ions across all relevant retention time windows of the corresponding precursor, allowing for accurate and comprehensive sample analysis. Precursor trails help compare fragmentation tails of different peptides and calculate similarity features to identify differences in fragmentation efficiency.

During database searching, LooMS extracts measures such as fragment ion count, complementary ions count, and consecutive ions count. Complementary ions are pairs formed from the same precursor ion, such as b ions (N-terminus product) and y ions (C-terminus product), which together help identify and characterize peptides. LooMS also extracts measures about the similarity of shape for matched fragment ions and between matched precursors and their fragments. Cosine correlations detect shape similarity to identify discrepancies and common patterns between precursors and fragments. Mass error, expressed in parts per million (ppm), measures the difference between the expected and actual ion mass. For each matching ion, mass errors are calculated by subtracting the theoretical mass from the measured mass. LooMS uses mass error measures to identify discrepancies between different PSMs. RT error measures the difference between the highest peak retention time of the precursor trail and the retention time of the ion with the smallest mass error.

LooMS scans for ion peaks with primary and secondary charges of b and y ions within the error tolerance range for the corresponding peptide across multiple retention time regions. Upon identifying the peptide, it accumulates values for ion count, complementary ions count, consecutive ions count, peak area, and peak half rank for b ions with primary and secondary charges, followed by y ions. When the precursor is undetectable, pseudo-precursors are generated to identify peptides with multiple charge states without an actual precursor peak signal. LooMS retains up to 10 corresponding peptides for each spectrum and extracts a total of 46 features for every PSM throughout different DIA data analysis stages.

#### 4. Retention Time Calibration Algorithm for PSM Data

After acquiring the PSM, LooMS uses a logistic regression model to analyze specific features and assign a score to the PSM. It then employs a retention time calibration algorithm, leveraging retention time data from the top 60,000 PSM records with the highest scores and predictions from the AutoRT tool. This algorithm uses dynamic programming to calibrate predicted retention times, excluding interference-matched PSMs outside the designated retention time range.

### 2.3. Samples separation

Traditional deep learning methods for scoring PSMs risk overfitting by using both target and decoy peptides, potentially leading to biased predictions. To address this, LooMS introduces a novel sample separation strategy. Instead of using a single decoy protein sequence, LooMS generates three decoy sequences based on the target protein sequences. This approach involves creating “open target” and “open decoy” protein sequences. Open target sequences combine the original target sequences with one decoy sequence, while open decoy sequences consist of the other two decoy sequences. LooMS then performs database searches using theoretical spectra from both open target and open decoy sequences against the DIA data. If a PSM originates from the open target sequence, which includes both target and decoy sequences, it is classified as a positive sample. Conversely, if the PSM arises from the open decoy sequence, it is categorized as a negative sample. Notably, the original label information of the sample is not used during model training for predicting the PSM score. This strategy helps mitigate the risk of overfitting, ensuring more robust and unbiased model performance.

### 2.4. Deep learning methods using iterative and bagging strategies

LooMS employs logistic regression scores to identify the top *N* samples, including the highest-scoring positive and negative samples for each experimental spectrum. These selected samples are used to train a deep learning model, which is then applied to score all PSMs. The training sample set is iteratively expanded through multiple iterations based on the current division of positive and negative samples. In each iteration, the highest-scoring negative samples and the corresponding positive samples within the top 200,000 are added to the training set. Positive samples with a FDR exceeding *K* % are filtered out to maintain sample quality. This iterative strategy enhances model accuracy by continually refining the training data and model parameters. Iterative strategies in deep learning involve a series of steps designed to improve model performance. These steps include training the model with different data sets, adjusting model parameters, and incorporating new samples. This approach helps reduce overfitting and improve model accuracy by identifying discrepancies between predictions and actual data, as well as uncovering patterns within the data.

## 3. Results

The experiment was conducted on a Linux server with a 2.50GHz Intel Xeon Gold 6248 CPU, Nvidia V100 16GB GPU, and 768GB of memory. Each method required specific input file formats; hence, the original .raw files were converted accordingly. Table 1 lists the parameter settings for LooMS and DiaNN (version 1.8.1) to ensure experimental objectivity.

**Table 1:** The parameters setting for DiaNN and LooMS.

|  | DiaNN | WaterloMS |
| --- | --- | --- |
| Human Dataset | 40856 | 57446 |
| Mouse Dataset | 98975 | 125307 |
| -fasta | 1 Decoy + 'R' + 1 Target | 3 Decoy and 1 Target |
| -qvalue | 0.01 | 0.01 |
| -min-fr-mz | 200 |  |
| -max-fr-mz | 1800 |  |
| -cut | K*,R*,!*P' | K*,R*,!*P' |
| -missed-cleavages | 2 | 2 |
| -min-pep-len | 6 | 6 |
| -max-pep-len | 98 | 98 |
| -min-pr-mz | 300 |  |
| -max-pr-mz | 1800 |  |
| -min-pr-charge | 1 | 1 |
| -max-pr-charge | 8 | 8 |

To evaluate the performance of LooMS, we applied it to human and mouse datasets with distinct gradients obtained from publicly available sources[24], accessible at ProteomeCentral. We used a 120-minute DIA gradient for the human dataset and a 240-minute DIA gradient for the mouse dataset, both commonly employed for analyzing complex protein mixtures. Reference protein sequences for each species were obtained from UniProt (March 2022), allowing for the generation of four protein sequences per species using targetdecoy strategies. Peptides were extracted from these sequences using trypsin with up to two missed cleavages. Spectral libraries were constructed, and retention time predictions were made using Prosit and AutoRT.

In the human dataset, LooMS detected 48,448 unique peptides (considering different charge states as a single entity) at a 1% FDR. In comparison, DiaNN identified 40,856 unique peptides at the same FDR level. In the mouse dataset, a parallel analysis demonstrates the presence of analogous outcomes. Specifically, LooMS identified 101,129 unique peptides, while DiaNN identified 98,975 unique peptides. The results are illustrated in Figure 2.

**Figure 2:**
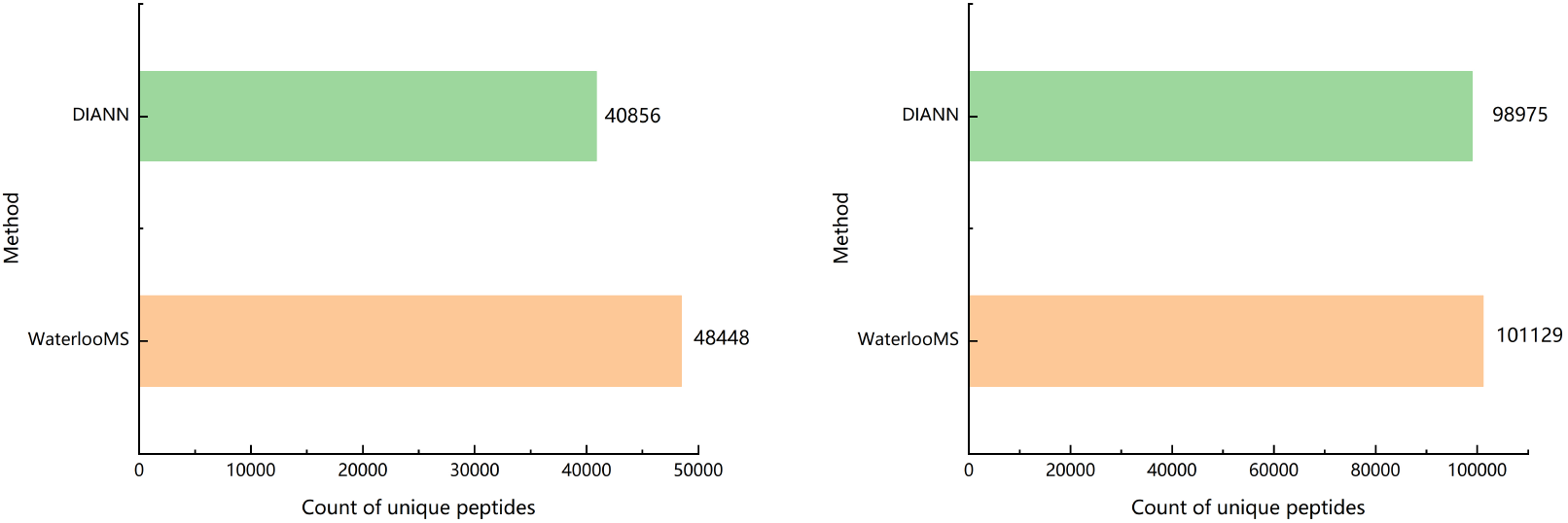
Comparison for count of unique peptides between LooMS and DiaNN. A. In human dataset, B. In mouse dataset.

Based on these findings, it is evident that LooMS exhibited a performance superiority of 18.58% compared to DiaNN when utilizing a 120-minute gradient. Moreover, LooMS marginally outperformed DiaNN by a slight margin in the context of a 240-minute gradient.

Interestingly, an analysis of the peptide overlap between the two methods at a 1% FDR demonstrated that 45% of the peptides identified by either method were common, as shown in figure 3.

**Figure 3:**
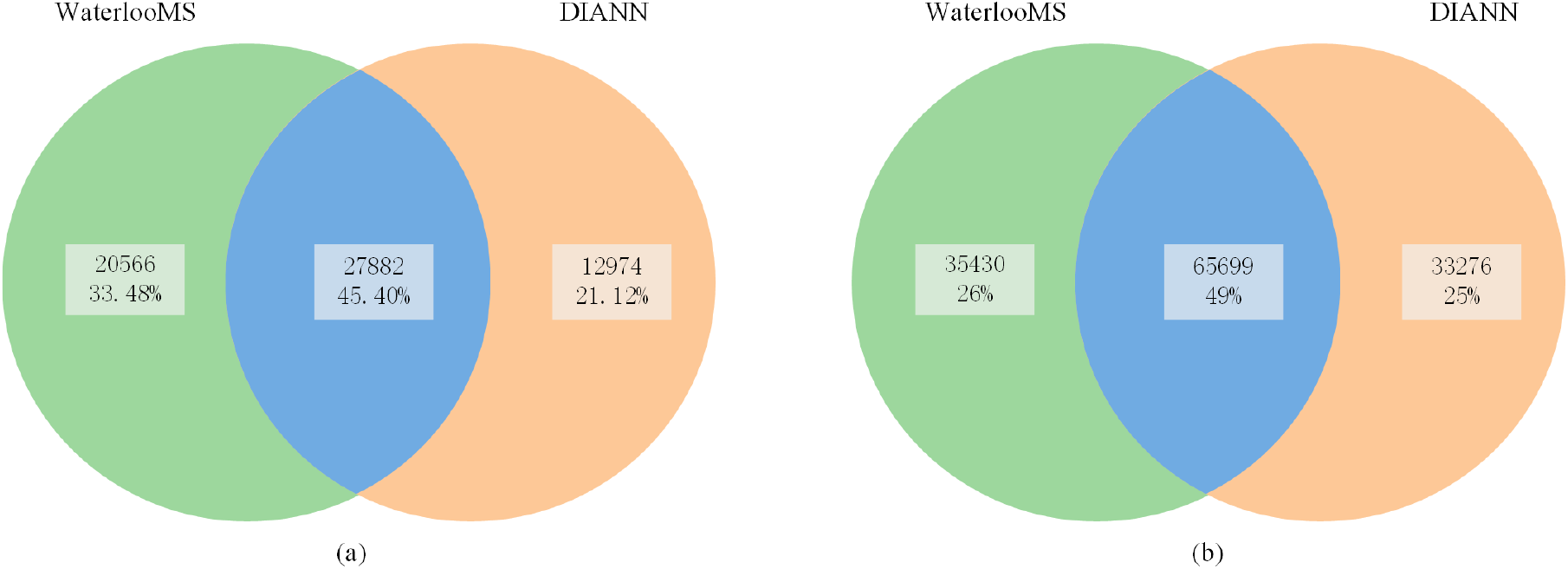
The overlapping of unique peptides identified by LooMS and DiaNN. A. In human dataset, B. In mouse dataset.

We observed that certain peptides classified as highly confident by one method (at a 1% FDR) also appeared in the results of the other method but did not meet the same threshold. This led us to integrate the results from LooMS and DiaNN using an innovative FDR control approach. Peptide scores from both methods were sorted in descending order, and initial threshold values *t1* and *t2* were set to 1. We selected the top *t1* peptides from LooMS and the top *t2* peptides from DiaNN, then calculated the ratio of decoy to target peptides for each method. If the ratio exceeded a specific threshold, *t1* and *t2* were updated to include more peptides. This process continued until all subsequent ratios exceeded the threshold. The overall FDR was then calculated, ensuring balanced integration of results. This method yielded significant improvements, with enhancements of 40.61% and 26.60% at the unique peptide level for the human and mouse datasets, respectively, as shown in Table 2.

**Table 2:** The comparison of unique peptides identified by DiaNN and method combined LooMS and DiaNN.

|  | # DiaNN | # Combine<br>WaterloMS with DiaNN | improvement rate |
| --- | --- | --- | --- |
| Human Dataset | 40856 | 57446 | 40.61% |
| Mouse Dataset | 98975 | 125307 | 26.60% |

## 4. Discussion

LooMS is a novel tool for identifying peptides in DIA datasets, utilizing an innovative strategy for unbiased positive and negative sample generation. This approach avoids the overfitting often associated with deep learning methods that use true labeled samples for training models to score PSMs during testing. Traditional methods rely on true target and decoy labels, training the model on a limited number of such samples even with transfer learning. LooMS generates positive samples by mixing target and decoy labels, preventing the model from directly accessing true label information. However, testing both LooMS and DiaNN with combined target and decoy protein sequences results in slower performance compared to using only the target protein sequence. This slowdown is due to the model ‘s prior knowledge of the label information, leading to overfitting.

During the process of model training, the selection of features is crucial. The predictive results of the model essentially provide a scoring function for measuring the degree of matching between experimental spectra and peptide segments. A good deep learning model should be able to comprehensively consider various factors related to experimental peak and peptide segment matching and properly integrate these factors to improve the accuracy and reliability of peptide detection in mass spectrometry. The present tool comprehensively considers various aspects of data analysis for DIA mass spectrometry and builds a large number of features for training deep learning models, which involve different stages of DIA data analysis. In the preprocessing stage of raw mass spectrometry data, the main focus is on obtaining quality control attributes of precursor and fragment ion trails. In the database search stage, the main considerations are the quality error of ions matched to theoretical amino acids, the number of consecutive ion matches, the number of complementary ion pairs, the abundance of ion peaks in the retention time region, the cosine similarity between the shape of precursor peaks and the shape of secondary spectrum peaks, and the cosine similarity between the shape of all matched secondary ion peaks. We also make full use of existing spectral library prediction tools and consider features such as the error of predicted retention time correction, the cosine similarity between the predicted secondary spectrum and the experimental spectrum, and the cosine similarity between the six highest-abundance matched precursor ion peaks and the predicted secondary spectrum peaks. The integration of the aforementioned features into the deep learning model resulted in a positive correlation between the peptide identification scores and the number of matched ion peaks, as well as the number of consecutive and complementary ion peaks. As the peptide identification score increased, the number of matched ion peaks, consecutive matches, and complementary matches also increased significantly. These findings suggest that the model is capable of providing a reliable scoring function that can effectively interpret mass spectrometry data when combined with intermediate results from the database search, thus increasing the reliability of peptide identification.

The integration of results from two tools can significantly improve the identification of peptides due to the differences in the features involved in peptide detection during their respective design. LooMS tool focuses mainly on the appearance of multiple consecutive retention time windows for the same precursor in the raw data. In the process of matching the secondary spectra and peptides, it introduces multiple consecutive time window signal data. LooMS considers the signal peak area, mass error, and retention time error formed by the trail during ion peak detection. On the other hand, DiaNN tool primarily focuses on the secondary signal peak of a specific retention time window during ion peak detection. The combination of results from both tools can synergistically utilize their strengths. A method was designed to control the FDR after merging the results by taking into account the PSM scores of both tools.

LooMS effectively leverages existing high-quality spectral library prediction tools, such as autoRT and Prosit, to generate features for multiple peptide detection models and improve their predictive performance. However, this approach also introduces certain limitations, such as the requirement to run these two spectral library software tools to obtain prediction results as input for LooMS when analyzing a new species ‘ sequence. Nevertheless, since the prediction results of autoRT and Prosit tools are invariant for the same peptide and parameters, this step need not be repeated for each peptide detection. The prediction results can be reused for the same sequence in a new species after performing spectral library prediction once.

## 5. Conclusion

LooMS represents an open-source, model-unbiased tool for predicting DIA peptides, which is released source code on GitHub at https://github.com/LooMS/dia_data_reading/tree/dia_integerated_looms. The tool effectively integrates features derived from multiple stages of peptide detection, demonstrating robust predictive performance. Notably, when combined with the widely recognized DiaNN tool, LooMS exhibits a substantial improvement exceeding 26%. Future research efforts will prioritize the integration of transfer learning models for predicting spectral library data.

## Notes

### Competing Interest Statement

The authors have declared no competing interest.

